# TORC1 phosphorylates and inhibits the ribosome preservation factor Stm1 to activate dormant ribosomes

**DOI:** 10.1101/2022.08.08.503151

**Authors:** Sunil Shetty, Jon Hofstetter, Stefania Battaglioni, Danilo Ritz, Michael N. Hall

## Abstract

Target of rapamycin complex 1 (TORC1) promotes biogenesis and inhibits degradation of ribosomes in response to nutrient availability. To ensure a basal supply of ribosomes, cells preserve a small pool of dormant ribosomes under nutrient-limited conditions. The regulation of dormant ribosomes is poorly characterized. Here, we show that upon inhibition of TORC1 by rapamycin or nitrogen starvation, Stm1 (suppressor of target of Myb protein 1) forms non-translating, dormant 80S ribosomes. Furthermore, Stm1-bound 80S ribosomes are protected from proteasomal degradation. Upon re-feeding, TORC1 directly phosphorylates and inhibits Stm1, thereby reactivating translation. Finally, SERBP1 (SERPINE1 mRNA binding protein), a mammalian ortholog of Stm1, forms dormant 80S ribosomes upon mTORC1 inhibition in mammalian cells. Thus, TORC1 regulates ribosomal dormancy in an evolutionarily conserved manner via a ribosome preservation factor.

## Introduction

The ribosome pool is tightly regulated to ensure appropriate translational capacity. Under nutrient-rich conditions, cells enhance ribosome biogenesis to promote protein synthesis. In contrast, under nutrient-limiting conditions, cells reduce ribosome biogenesis and degrade ribosomes via autophagy (ribophagy) or proteasomal degradation (An & Harper, 2020; Kraft *et al*, 2008; Prossliner *et al*, 2018; Wang *et al*, 2020). However, excessive degradation may deplete ribosomes and impede resumption of cell growth upon restoration of favorable growth conditions (Kressler *et al*, 2010; Warner, 1999). To avoid excessive degradation during starvation, so-called preservation factors protect a small pool of non-translating, vacant ribosomes (Ben-Shem *et al*, 2011; Brown *et al*, 2018; Prossliner *et al*., 2018; Wells *et al*, 2020). The regulation of the preservation and subsequent re-activation of mRNA-free dormant ribosomes is not yet understood.

TORC1 is an evolutionarily conserved serine/threonine kinase that promotes cell growth in response to nutrients and growth factors (Battaglioni *et al*, 2022; Gonzalez & Hall, 2017; Saxton & Sabatini, 2017). In budding yeast, TORC1 comprises TOR1/TOR2, Kog1, and Lst8 while mammalian TORC1 (mTORC1) consists of mTOR, the Kog1 ortholog RAPTOR, and mLST8. Under nutrient-sufficient conditions, TORC1 up-regulates ribosome biogenesis by promoting transcription of rRNAs and expression of ribosomal proteins, and prevents ribosomal degradation by inhibiting ribophagy and the proteasome (An & Harper, 2020; Kraft *et al*., 2008; Mayer & Grummt, 2006). Inhibition of TORC1 by starvation or rapamycin treatment reduces translation and leads to the accumulation of 80S ribosomes (Barbet *et al*, 1996; Larsson *et al*, 2012). The accumulation of 80S ribosomes rather than free 40S and 60S subunits upon TORC1 inhibition is unexplained.

Stm1 is a yeast ribosome preservation factor (Van Dyke et al., 2013). In vacant 80S ribosomes isolated from starved cells, Stm1 is at the interface of the two ribosomal subunits and occupies the mRNA tunnel of the 40S subunit, thereby excluding mRNA (Ben-Shem et al., 2011). Stm1 was also shown to interact with translating ribosomes (Van Dyke et al., 2006; Van Dyke et al., 2009). Furthermore, cells lacking Stm1 are hypersensitive to nitrogen starvation or rapamycin treatment (Van Dyke et al., 2006), suggesting a functional but so far poorly characterized interaction between Stm1 and TORC1.

Here we show that upon TORC1 inhibition or nitrogen starvation, Stm1 forms vacant 80S ribosomes. Furthermore, Stm1-bound 80S ribosomes are protected from proteasomal degradation. Upon restoration of growth conditions, TORC1 directly phosphorylates and inhibits Stm1, thereby activating dormant ribosomes and stimulating translation re-initiation. Such regulation of dormant ribosomes is evolutionarily conserved as mTORC1 similarly regulates SERBP1 in mammalian cells.

## Results

### Stm1 occupies 80S ribosomes upon TORC1 inhibition

TORC1 inhibition or nutrient depletion leads to accumulation of non-translating 80S ribosomes (Ashe *et al*, 2000; Barbet *et al*., 1996; Kaminskas, 1972; Liu & Qian, 2016; van Venrooij *et al*, 1972). Why 80S ribosomes rather than free 40S and 60S subunits accumulate is not understood. To investigate this, we performed mass spectrometry on 80S ribosomes isolated from rapamycin-treated and -untreated cells (Figure 1A). We observed enrichment of factors involved in ribosome preservation, translation initiation, ribosome biogenesis, or mRNA degradation in the 80S fraction from rapamycin-treated cells compared to -untreated cells. These factors included Stm1, Dbp2, Mrt4, Asc1, Ski2, Sui3, and Ssb1 (Figure S1A). Stm1 was particularly prominent among the proteins enriched in the 80S fraction from TORC1-inhibited cells (Figures 1A and 1B).

**Figure 1.**
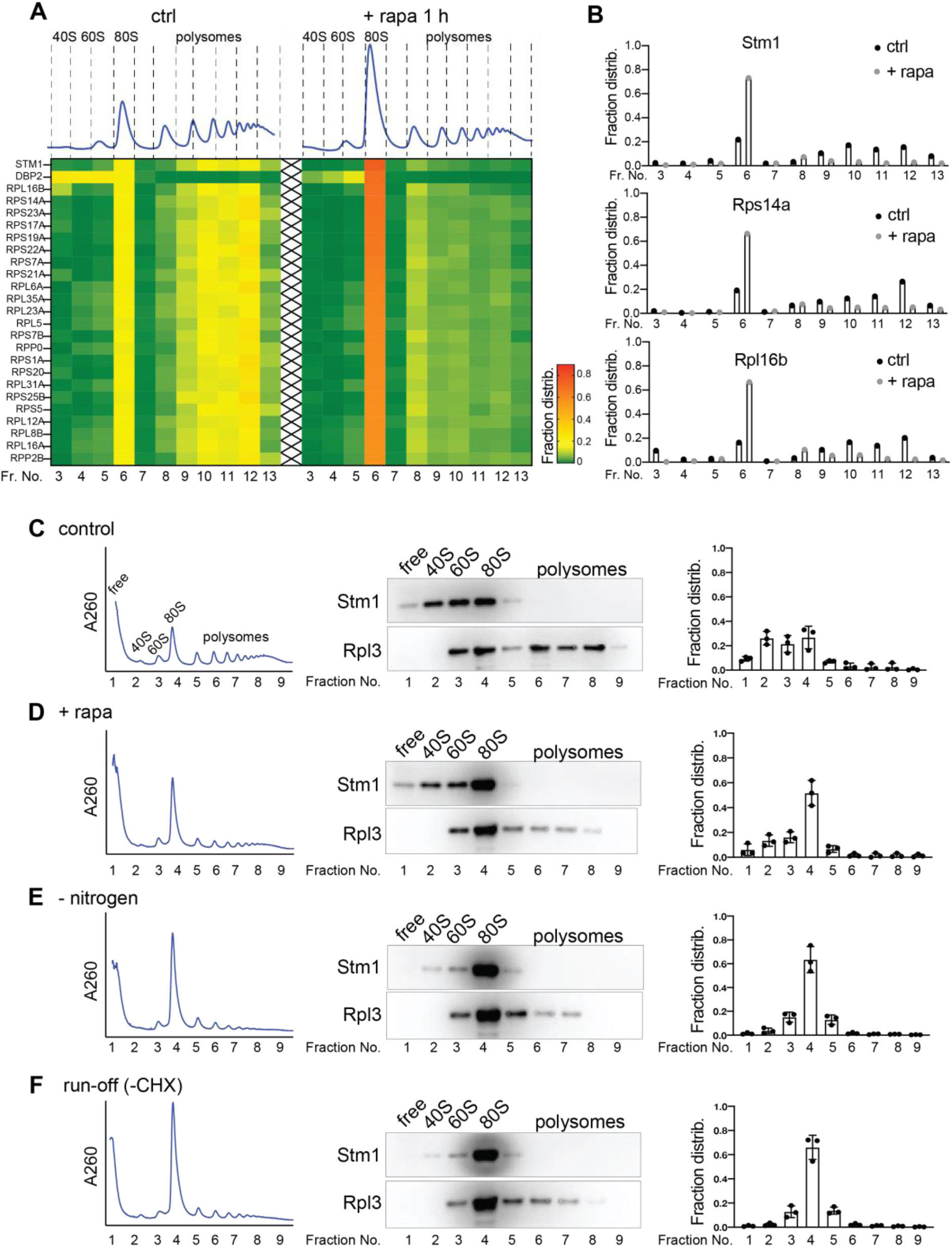
Stm1 binds to vacant 80S ribosome upon TORC1 inhibition. **A.** Heat map showing the fraction distribution of the top 25 most abundant proteins across the polysome profiles treated with or without rapamycin for 1 hour. The proteins are sorted according to their distribution in the 80S fraction (fraction 6) upon rapamycin treatment (Fraction number 6). Individual polysome profiles were separated on a 5-50% sucrose density gradient containing 50 mM KCl into 14 fractions, and fraction numbers 3 to 13 were analyzed by mass spectrometry. **B.** Distribution of proteins Stm1, Rps14a, and Rpl6b in the fractions across the polysome profile obtained with or without rapamycin treatment for 1 hour as identified by MS/MS. **C-F**. Polysome profiling of yeast cells with (**C-E**) or without (**F**) cycloheximide (CHX) treatment, immunoblots for Stm1 and Rpl3 in polysome fractions, and quantification of Stm1 distribution across the polysome profiles of cells grown under nutrient-rich condition (**C, F**), rapamycin treatment for 1 hour (**D**), or nitrogen starvation for 1 hour (**E**). Individual polysome profiles were fractionated on a 5-50% sucrose density gradient containing 150 mM KCl into 9 fractions, precipitated and immunoblotted for Stm1 and Rpl3. Rpl3 is used as a loading control. Three biological replicates are used for the quantification of fraction distribution.

In the crystal structure of 80S ribosomes, Stm1 was found to occupy the mRNA tunnel in otherwise vacant 80S ribosomes, rendering them translationally inactive (Ben-Shem *et al*., 2011). However, Van Dyke *et al*. (2006) showed, and we confirmed (Figure S1B), that Stm1 associates with both 80S and polysomal ribosomes. This raises the question of how Stm1 is associated with translating polysomes? We speculated that Stm1 might associate non-specifically with polysomes, possibly due to the low KCl concentration (50 mM) in the sucrose density gradients of the above experiments. Non-specific interactions of proteins with ribosomes are known to be KCl sensitive (Fleischer *et al*, 2006). Hence, we re-analyzed the interaction of Stm1 with ribosomes using sucrose density gradients containing higher KCl concentrations (150 and 300 mM). Notably, at 150 mM KCl, the association of Stm1 with polysomes was lost whereas the interaction with 80S ribosomes was unaffected, suggesting that the interaction of Stm1 with polysomes is non-specific. We note that, at 150 mM KCl, Stm1 was also detected associated with the 40S and 60S subunits (Figures 1C and S1C). Furthermore, inhibition of TORC1 by rapamycin or nitrogen starvation led to an accumulation of Stm1-bound 80S ribosomes (Figures 1D, 1E, and S1F). To confirm that Stm1 is associated with non-translating 80S ribosomes, we analyzed polysome profiles in the absence of cycloheximide (-CHX), a condition that allows actively translating ribosomes to run-off mRNA. Ribosome run-off increased (∼ 3-fold) the accumulation of Stm1 in the 80S ribosome fraction compared to the CHX-treated samples (Figure 1F and S1F). Thus, Stm1 is associated with non-translating, dormant 80S ribosomes upon TORC1 inhibition or nutrient deprivation.

### Stm1 forms dormant 80S ribosomes upon TORC1 inhibition

To understand the role of Stm1 in vacant 80S ribosomes, we analyzed polysome profiles of wild-type and *stm1*Δ strains. We observed little difference in the overall polysome profiles of wild-type and *stm1*Δ cells grown in nutrient-rich conditions (Figure 2A), suggesting that Stm1 has no role in normally growing cells. Upon TORC1 inhibition by rapamycin treatment or nitrogen starvation for 1 hour, wild-type cells displayed the expected shift of polysomes to 80S ribosomes (Figures 2B and 2C). In contrast, *stm1*Δ cells displayed a shift of polysomes to 40S and 60S subunits, suggesting that Stm1 forms non-translating 80S ribosomes (Figures 2B and 2C). Confirming that Stm1 is required for the formation of vacant 80S ribosomes, ribosome run-off (polysome profiling in the absence of cycloheximide) led to increased accumulation of 40S and 60S subunits and a corresponding decrease in 80S ribosomes in *stm1*Δ cells grown in nutrient-rich conditions, compared to wild-type cells (Figure 2D).

**Figure 2.**
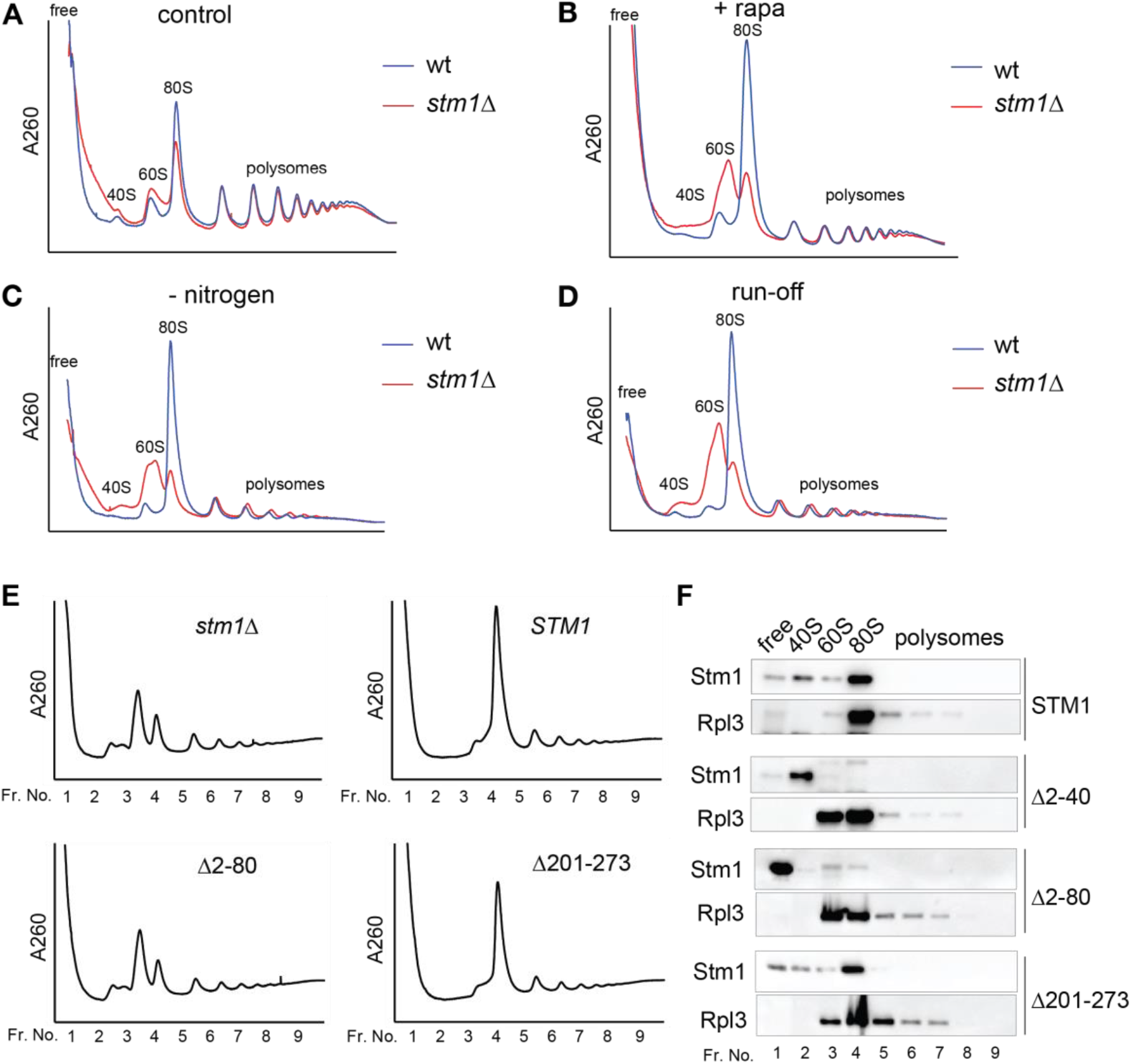
Stm1 forms vacant 80S ribosomes upon TORC1 inhibition. **A-D**. Polysome profiles of wild-type (wt) and *stm1*Δ cells with (**A-C**) or without (**D**) CHX treatment under nutrient-rich conditions (**A, D**), with rapamycin treatment for 1 hour (**B**), or under nitrogen starvation for 1 hour (**C**) using 5-50% sucrose density gradients containing 150 mM KCl. At least three biological replicates are analyzed for each condition. **E.** Polysome profiles of *stm1*Δ cells containing either full-length or N-terminally deleted or C-terminally deleted Stm1 mutants upon nitrogen starvation for 1 hour. **F**. Binding of full-length or deletion mutants of Stm1 across the polysome profile fractions shown in Figure 2E. 9 equal volume fractions of each polysome profile were precipitated and analyzed by immunoblot for Stm1 and Rpl3. At least two biological replicates are used for each mutant.

To determine which part of the Stm1 protein is required for the formation of vacant ribosomes, we generated a series of deletions (non-overlapping deletion of 40-80 amino acids) covering the entire Stm1 protein (Figure S2A). C-terminal deletion mutants of Stm1 (lacking amino acids 161-200, 201-240 or the C-terminal residues 201-273) formed 80S ribosomes upon nitrogen starvation, similar to wild-type Stm1 (Figures 2E, 2F, S2B, and S2C). However, Stm1 mutants lacking N-terminal regions (lacking amino acids 2-40, 2-80, 41-80, 81-120) failed to form 80S ribosomes upon nitrogen starvation (Figures 2E, 2F, S2B, and S2C). According to the crystal structure of Stm1-bound 80S ribosomes, residues 9 to 40 of Stm1 interact with the 60S subunit while residues 60 to 141 associate with the 40S subunit (Ben-Shem *et al*., 2011). Amino acids 40 to 60 of Stm1 act as a linker between the two ribosomal subunits. Thus, the N-terminal region of Stm1 is required for the formation of dormant 80S ribosomes.

Stm1 was reported to be important for survival in the presence of rapamycin (Van Dyke *et al*, 2006). Consistent with the previous reports, we also observed that deletion and overexpression of Stm1 confer rapamycin hypersensitivity and resistance, respectively (Figures S2D and S2E). Stm1 mutants lacking their N-terminal region could not completely rescue the rapamycin hypersensitivity of *stm1*Δ cells while C-terminal mutants could (Figure S2F). Thus, the N-terminal region of Stm1, which is involved in dormant ribosome formation, is important for coping with stress upon TORC1 inhibition.

### TORC1 phosphorylates and inhibits Stm1

How does TORC1 control the interaction of Stm1 with 80S ribosomes? To determine if Stm1 is regulated by TORC1 via phosphorylation, we performed a phosphoproteomic analysis of total extracts and immunoprecipitated Stm1-GFP from rapamycin-treated and - untreated yeast cells (Figure S3A and S3B). This revealed that several residues in Stm1 are phosphorylated, of which S32, S41, S45, S55, S73, T181, and T218 were significantly less phosphorylated upon rapamycin treatment. Interestingly, in the crystal structure of Stm1-bound 80S ribosomes, serines 41, 45, and 55 are in the Stm1 linker region spanning the space between the 40S and 60S subunits (Ben-Shem *et al*., 2011). Furthermore, these three serines fall within the N-terminal region of Stm1 that mediates formation of dormant 80S ribosomes (see above). Hence, we hypothesize that phosphorylation at these sites plays a particularly important role in the regulation of Stm1.

To assess whether TORC1 directly phosphorylates Stm1 on these three serine residues, we performed an in vitro kinase assay using recombinant mammalian TORC1 (mTORC1) and Stm1. mTORC1 phosphorylated wild-type Stm1 (Figure 3A). Serine-to-alanine mutation of the three serines (Stm1^AAA^, S41A/S45A/S55A) reduced the mTORC1-mediated phosphorylation of Stm1 (Figure 3A and 3B). We note that the Stm1^AAA^ mutant could still be phosphorylated by mTORC1, likely due to phosphorylation of the other serine residues identified in our phosphoproteomic analysis (Figure S3A). Thus, Stm1 is a novel TORC1 substrate.

**Figure 3.**
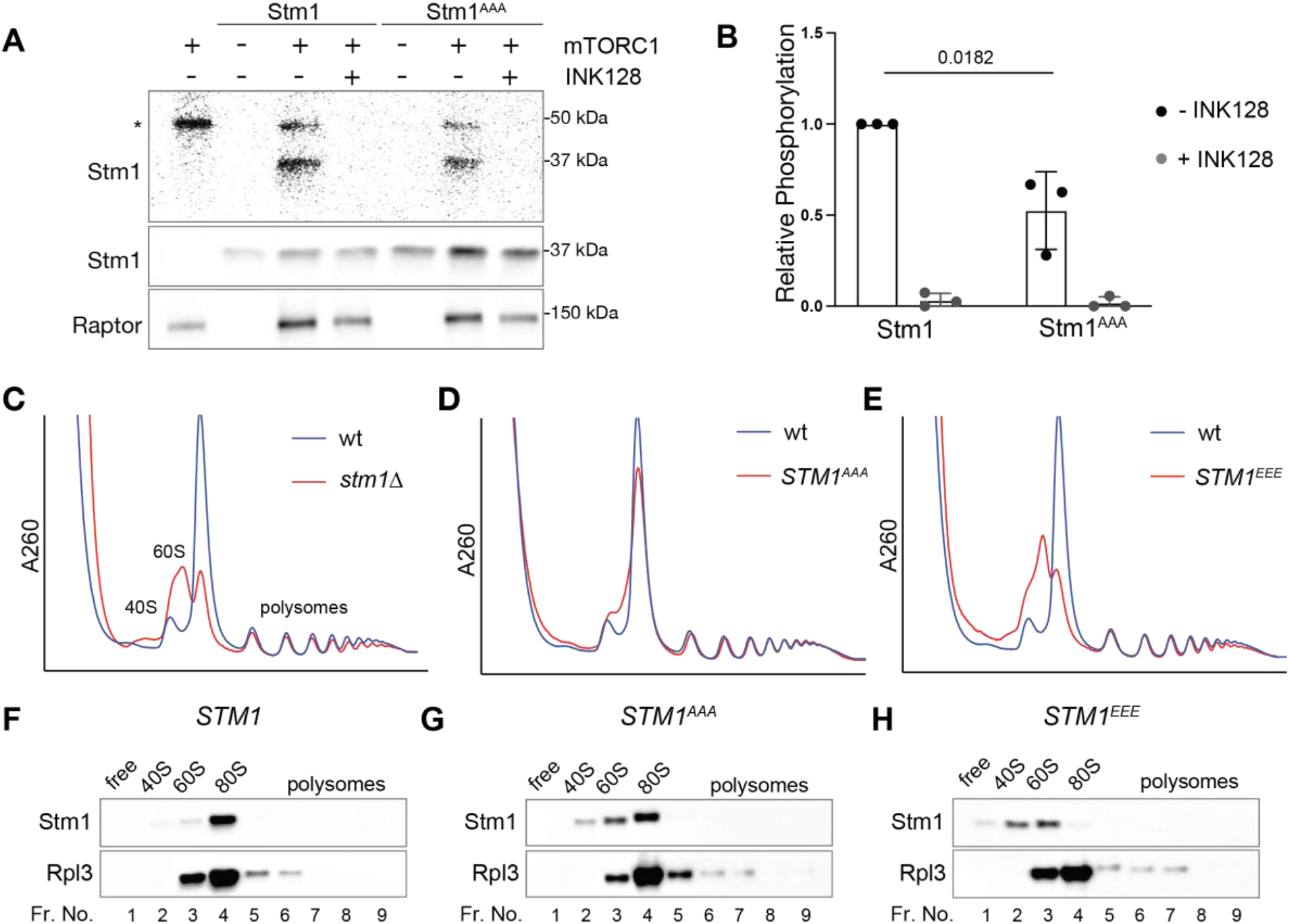
TORC1 phosphorylates and inhibits Stm1. **A-B**. In vitro kinase assay (**A**) and its quantitation (**B**) using recombinant mTORC1 and recombinant Stm1 or Stm1^AAA^ as a substrate in the presence of ^32^P-γ-ATP. Immunoblots for Stm1 and Raptor are shown. Asterisk indicates the auto-phosphorylation of a mTORC1 component. INK128 is used as an inhibitor of mTOR. Minimum of three replicates are included and P-value is calculated based on paired t-test. **C.** Polysome profiles of wild-type and *stm1*Δ cells under nitrogen starvation for 1 hour. **D.** Polysome profiles of wild-type and *STM1^AAA^* cells under nitrogen starvation for 1 hour. **E.** Polysome profiles of wild-type and *STM1^EEE^* cells under nitrogen starvation for 1 hour. **F-H.** Immunoblots for Stm1 and Rpl3 from the ribosomal fractions obtained from wild-type (**F**), *STM1^AAA^*(**G**), and *STM1^EEE^* (**H**) cells under nitrogen starvation for 1 hour.

To elucidate the role of TORC1-mediated Stm1 phosphorylation in the formation of dormant ribosomes, we analyzed polysome profiles of *stm1*Δ cells expressing either phospho-deficient (Stm1^AAA^) or a phospho-mimetic (Stm1^EEE^, S41E/S45E/S55E) Stm1. Rapamycin-treated *STM1*^AAA^ cells formed 80S ribosomes like wild-type cells (Figures 3D, 3F, and 3G). However, *STM1*^EEE^ cells, like *stm1*Δ cells, accumulated the 40S and 60S subunits and were thus defective in forming 80S ribosomes, upon rapamycin treatment (Figures 3C, 3E, and 3H). These results suggest that TORC1-mediated phosphorylation of at least the linker region of Stm1 inhibits the formation of dormant, Stm1-bound 80S ribosomes.

### Stm1 protects ribosomes from proteasomal degradation

Stm1, as a preservation factor, was reported to prevent ribosome degradation in long-term stationary phase cells (Van Dyke *et al*, 2013). Thus, we examined the effect of long-term rapamycin treatment (24 hours) on the ribosome content of cells in which Stm1 expression is altered. Deletion or overexpression of Stm1 had no effect on ribosome levels in the absence of rapamycin, compared to wild-type (Figures S4A). In the presence of rapamycin, *stm1*Δ cells contained fewer total 40S and 60S subunits, compared to control cells (Figure S4B). Cells overexpressing Stm1 contained more ribosomes compared to control cells (Figure S4B). Importantly, the level of ribosomes in different conditions was confirmed by immunoblotting for Rpl3 (Figure S4C). These findings support the observation of Van Dyke et al. (2013) that Stm1 is a ribosome preservation factor.

Next, we investigated the role of TORC1-mediated Stm1 phosphorylation on ribosome preservation. Similar to the *stm1*Δ, *STM1^EEE^* cells exhibited lower ribosome content compared to wild-type or *STM1^AAA^* cells, upon long-term rapamycin treatment (Figures 4A and 4B). *stm1*Δ and *STM1^EEE^* cells also showed lower ribosome content upon long-term (24 hours) nitrogen starvation (Figures S4D). Immunoblot analysis also showed reduced levels of ribosomal protein Rpl3, in both *stm1*Δ and *STM1^EEE^*cells, upon rapamycin treatment (Figures 4C and 4D) or nitrogen starvation (Figures S4E and S4F). Analysis of polysome profiles and Rpl3 levels of normally growing cells revealed no change in ribosome content, regardless of the status of Stm1 (Figures S4G, S4H, and S4I). These data suggest that dephosphorylation of Stm1 upon TORC1 inhibition, and Stm1’s subsequent formation of dormant 80S ribosomes, are important for the long-term preservation of ribosomes.

**Figure 4.**
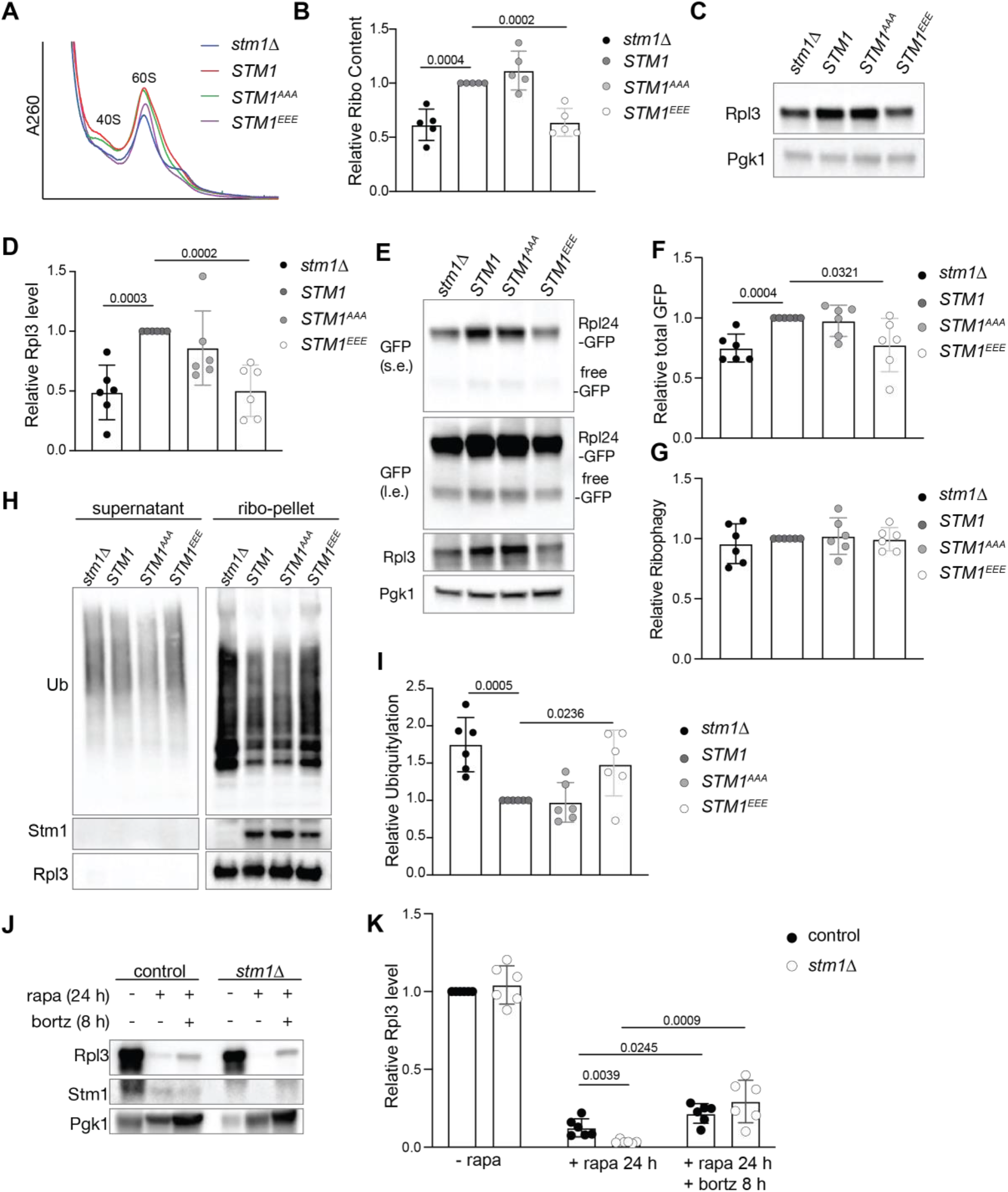
Stm1 protects ribosomes from proteasomal degradation. **A-B.** Ribosomal subunit profiles (**A**) and quantitation of ribosome content (**B**) of *stm1*Δ, wild-type, *STM1^AAA^*, and *STM1^EEE^* cells treated with rapamycin for 24 hours. For the ribosomal subunit profile analyses, the cell extracts were treated with 50 mM EDTA and separated on 5-25% sucrose density gradients. The P-value from multiple t-tests is indicated above in the graphs. **C-D**. Immunoblot (**C**) and the quantification (**D**) of Rpl3 in *stm1*Δ, wild-type, *STM1^AAA^*, and *STM1^EEE^* cells treated with rapamycin for 24 hours. At least six biological replicates are subjected to statistical analysis using multiple t-tests. **E-G**. Immunoblot of Rpl24-GFP and free GFP (**E**), quantification of total GFP relative to Pgk1 (**F**), and free GFP relative to Rpl24-GFP (**G**) in *stm1*Δ, wild-type, *STM1^AAA^*, and *STM1^EEE^* cells treated with rapamycin for 24 hours. At least six biological replicates are subjected to statistical analysis using multiple t-tests. (s.e. for short exposure; l.e. for long exposure). **H-I**. Immunoblot for ubiquitin in the supernatant and ribosomal pellet (**H**) obtained from *stm1*Δ, wild-type, *STM1^AAA^*, and *STM1^EEE^* cells treated with rapamycin for 24 hours. Immunoblots for Stm1 and Rpl3 are shown as controls. Quantification of ribosomal ubiquitylation (**I**) relative to wild-type is shown. The ribosome pellets were obtained using a 1 M sucrose cushion. At least six biological replicates are subjected to statistical analysis using multiple t-tests. **J-K**. Immunoblot (**J**) and the quantification (**K**) of Rpl3 in wild-type and *stm1*Δ cells without rapamycin, or with rapamycin for 24 hours along with or without bortezomib treatment for 8 hours. At least six biological replicates are subjected to statistical analysis using multiple t-tests.

In yeast, upon TORC1 inhibition, ribosomes are degraded mainly via ribophagy (Kraft *et al*., 2008). Thus, we tested whether ribosomes in rapamycin-treated *stm1Δ* cells undergo increased degradation via ribophagy. In yeast, ribophagy can be monitored by tagging ribosomal proteins with a green fluorescent protein (GFP) and assaying the accumulation of free GFP which is resistant to vacuolar proteases (Kraft *et al*., 2008). Upon long-term rapamycin treatment (24 hours), *stm1*Δ and *STM1^EEE^* cells contained less Rpl24-GFP and, surprisingly, also less free GFP, compared to control cells (Figures 4E, 4F, S4J, and S4K). However, there was no difference in the ratio of free GFP to Rpl24-GFP in wild-type, *stm1*Δ, and *STM1^EEE^*cells (Figures 4G and S4L), suggesting that the enhanced degradation of ribosomes in rapamycin-treated *stm1*Δ cells is not due to ribophagy.

An alternative pathway for ribosomal degradation is the ubiquitin-proteasome pathway (An & Harper, 2020). Indeed, analysis of poly-ubiquitin in isolated ribosomes from long-term rapamycin-treated cells revealed increased ribosome poly-ubiquitylation in *stm1Δ* cells compared to controls (Figures 4H, and 4I). *STM1^EEE^* cells, but not *STM1^AAA^* cells, also showed increased ribosome poly-ubiquitylation upon long-term rapamycin treatment (Figures 4H, and 4I). These data suggest that ribosomes are subjected to increased ubiquitylation in the absence of dormancy, following TORC1 inhibition. To rule out that the increased ubiquitylation of ribosomes is due to ubiquitylation of nascent polypeptides, we treated cells with puromycin for 15 min before harvesting to release nascent polypeptides. The puromycin treatment did not reduce the amount of ubiquitin in ribosomal pellets, further supporting increased poly-ubiquitylation of ribosomes in *stm1*Δ cells (Figure S4M). To investigate further whether ribosomes are degraded by the proteasome during rapamycin treatment, we treated cells with the proteasome inhibitor bortezomib. This experiment was performed in a strain lacking Pdr5 (a multidrug efflux pump) to make yeast cells responsive to bortezomib (Collins *et al*, 2010). GFP pull-down of ribosomes using Rpl24-GFP revealed increased ribosomal ubiquitylation in *stm1Δ* and *STM1^EEE^* cells upon long-term rapamycin treatment (Figures S4N and S4O). Bortezomib further enhanced the poly-ubiquitylation of ribosomes in *stm1Δ* and *STM1^EEE^*cells. Furthermore, bortezomib increased Rpl3 levels in rapamycin-treated *stm1Δ* and wild-type cells (Figures 4J and 4K). These results suggest that Stm1, in addition to mediating the formation of dormant ribosomes, preserves 80S ribosomes by preventing their ubiquitylation and proteasomal degradation.

### TORC1 activates dormant ribosomes by inhibiting Stm1

The above findings suggest that cells maintain a pool of dormant ribosomes during long-term starvation, possibly to kick-start growth upon re-feeding. Similar to a previous study (Van Dyke *et al*., 2013), we observed that resumption of translation upon re-feeding of starved cells was delayed in *stm1*Δ cells (Figure S5A). Thus, Stm1-mediated preservation of ribosomes under starvation is required for efficient translation recovery upon re-feeding. To investigate whether TORC1-mediated phosphorylation of Stm1 promotes resumption of translation, we analyzed translation in re-fed wild-type, *STM1^AAA^*, or *STM1^EEE^* mutant cells. Wild-type cells exhibited recovery of translation within 10 minutes of nutrient replenishment, while both *STM1^AAA^*and *STM1^EEE^* cells required >30 minutes for resumption of translation, as measured by puromycin incorporation (Figures 5A and 5B). Delayed recovery of translation in *STM1^EEE^* and *stm1*Δ cells correlated with reduced ribosome content (Figures S4D and S4E). Importantly, *STM1^AAA^* cells displayed delayed recovery of translation despite normal ribosome levels (Figures S4D and S4E). Under nutrient-sufficient conditions, manipulation of Stm1 did not affect translation (Figures S5B and S5C). Altogether, this suggests that TORC1-mediated phosphorylation of Stm1 at serines 41, 45, and 55 is required for the efficient activation of dormant ribosomes and stimulation of translation during recovery from starvation.

**Figure 5.**
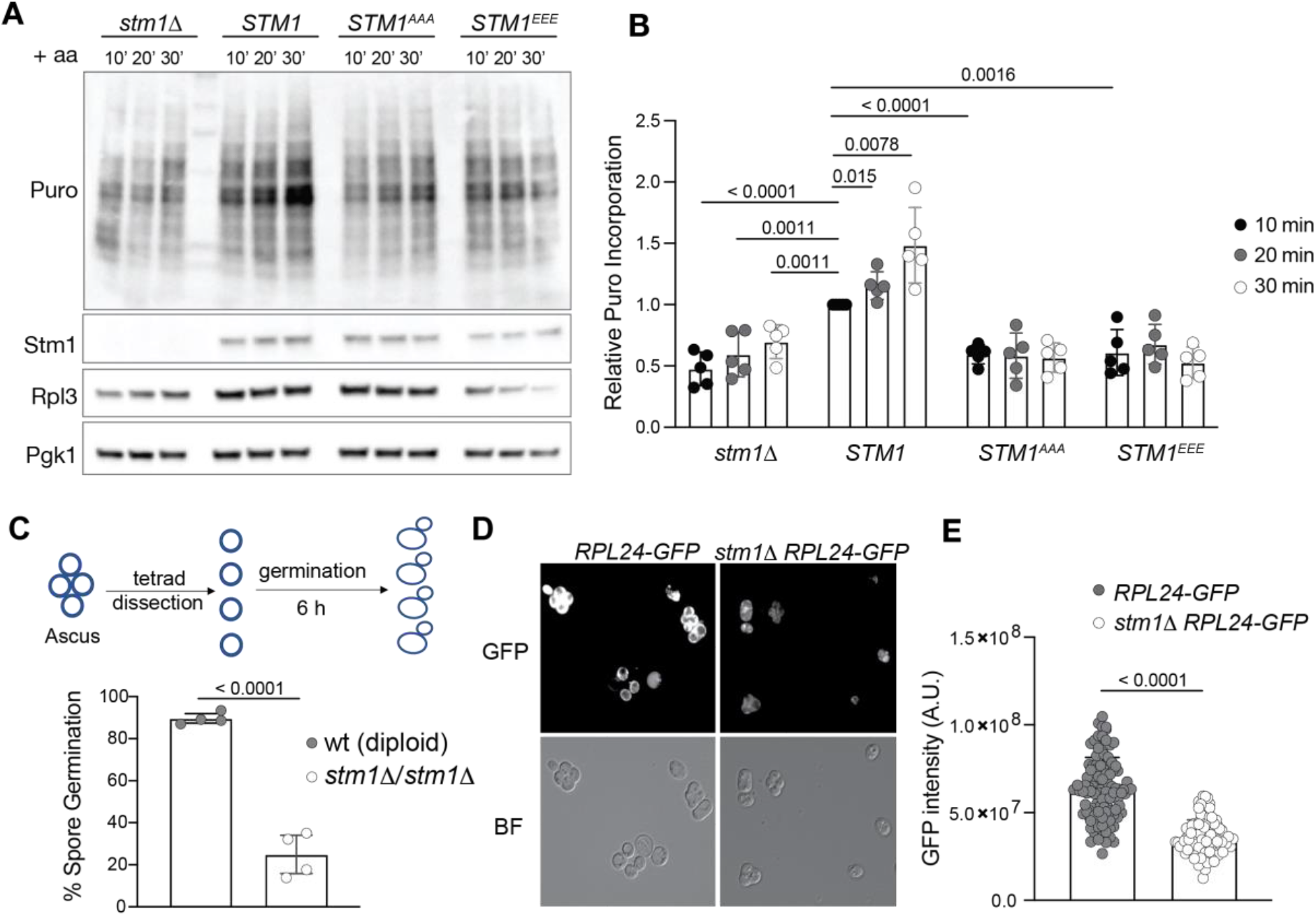
TORC1 activates dormant ribosomes by inhibiting Stm1. **A-B.** Puromycin incorporation (**A**) and its quantification (**B**) in *stm1*Δ, wild-type, *STM1^AAA^*, and *STM1^EEE^* cells starved of nitrogen for 24 hours and restimulated with amino acids for 10, 20, or 30 minutes. Immunoblots for puromycin, Stm1, Rpl3, and Pgk1 are shown. Relative puromycin incorporation for at least five replicates is subjected to multiple t-tests. **C**. Germination efficiency of spores upon dissection of tetrads (schematic in upper panel) from diploid wild-type and *stm1*Δ cells on YPD agar media after 6 hours of incubation. Minimum of four replicates are used for multiple t-tests. **D-E**. Representative fluorescent microscopic images for expression of Rpl24-GFP (**D**) and its intensity quantification (**E**) in tetrads of control (*RPL24GFP*) and *stm1*Δ *RPL24GFP* cells. Experiments are performed at least three times.

We investigated the significance of Stm1-mediated ribosome preservation in another physiological context. In response to long-term nitrogen and glucose starvation, diploid yeast cells undergo meiosis and form asci containing four haploid spores, equivalent to gametes, which then germinate upon nutrient replenishment. We examined the role of the Stm1-mediated preservation of ribosomes in spore germination. In particular, we examined the germination of spores derived from *stm1*Δ/*stm1*Δ diploid cells (Figure 5C). 90% of wild-type spores but only 25% of *stm1*Δ spores germinated within 6 hours of nutrient stimulation (Figure 5C). This was not due to the reduced viability of *stm1Δ* spores as essentially all spores eventually germinated to form small colonies after 2 days of incubation (Figures S5D and S5E). The level of ribosomal protein Rpl3 was decreased in spores derived from *stm1*Δ/*stm1*Δ cells, compared to spores from wild-type diploid cells (Figure S5F), suggesting a loss of ribosomes in *stm1*Δ spores. To confirm a reduction in ribosomes in *stm1Δ* spores, we monitored the level of GFP-tagged Rpl24 in spores from *stm1*Δ/*stm1*Δ diploid cells. The level of Rpl24-GFP was significantly decreased in *stm1Δ* spores, compared to wild-type spores (Figures 5D, 5E, S5G, and S5H). These findings suggest that Stm1-mediated preservation of ribosomes is required for efficient germination of dormant spores.

### Regulation of ribosome dormancy by TORC1 is evolutionarily conserved

The interaction of Stm1 with ribosomes appears to be conserved in eukaryotes as the structure of vacant human 80S ribosomes contained SerpineE1 mRNA binding protein (SERBP1), an ortholog of yeast Stm1 (Anger *et al*, 2013; Brown *et al*., 2018). We investigated whether mTORC1 affects the interaction of SERBP1 with 80S ribosomes. In polysome profiles from proliferating HEK293T cells, SERBP1 was found preferentially in the 40S fraction but also in the 60S, 80S, and low MW polysome fractions (Figure 6A). Inhibition of mTOR with INK128 caused SERBP1 to accumulate in the 80S fraction (Figures 6B and S6A). Increasing the amount of 80S ribosomes by harringtonine treatment, which traps mRNA-bound 80S initiation complexes, did not increase the amount of SERBP1 in the 80S fraction (Figure 6C). These results suggest that SERBP1, like Stm1, forms a complex only with mRNA-free 80S ribosomes. A previous study reported an interaction of SERBP1 mainly with translating ribosomes (polysomes) (Muto *et al*, 2018). As observed with Stm1 (see above), the interaction of SERBP1 with polysomes was dependent on the concentration of KCl (50 vs. 150 mM), again suggesting the interaction was non-specific (Figures S6A and S6B).

**Figure 6.**
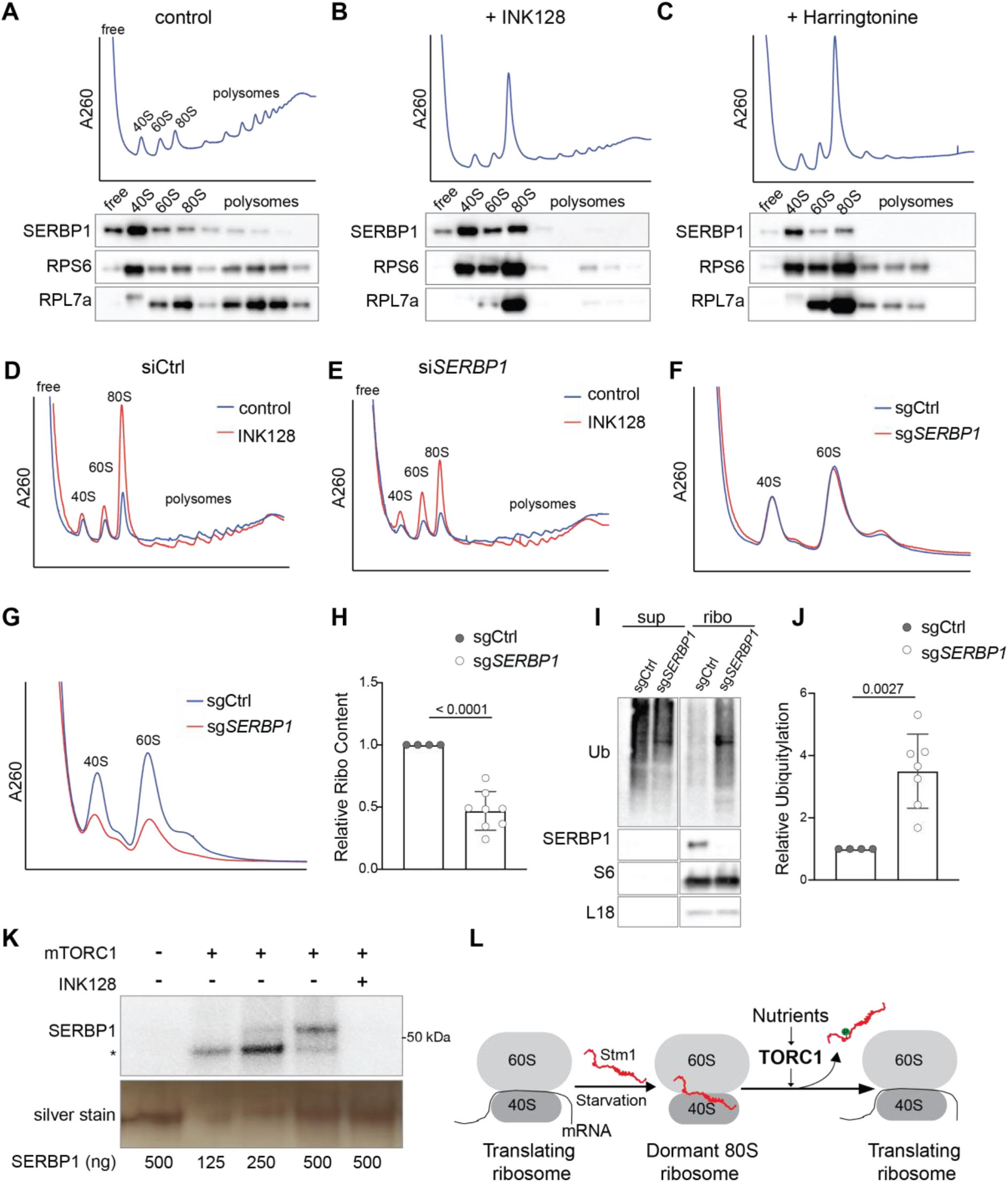
SERBP1 preserves ribosomes in human cells. **A-C.** Polysome profile and immunoblotting of SERBP1, RPS6, and RPL7a across the polysome fractions of HEK293T cells treated with DMSO (**A**), INK128 (**B**), or harringtonine (**C**) for 1 hour. Polysome profiles were performed using 5-50% sucrose density gradients containing 150 mM KCl. **D-E**. Polysome profile analyses of HEK293T cells treated with control siRNA (siCtrl) (**D**) or SERBP1 siRNA (siSERBP1) (**E**), with or without INK128 treatment for 1 hour. Polysome profiles were performed using 5-50% sucrose density gradients containing 150 mM KCl. **F.** Ribosome subunit analysis of HEK293T cells treated with control sgRNA (sgCtrl) or SERBP1 sgRNA (sgSERBP1) under normal conditions. Minimum of four replicates are analyzed. **G-H**. Ribosome subunit analysis (**F**) and its quantification (**G**) of HEK293T cells treated with control sgRNA (sgCtrl) or SERBP1 sgRNA (sgSERBP1) along with INK128 treatment for 48 hours. **I-J**. Immunoblot for ubiquitin (**I**) and its quantification (**J**) in the supernatant and ribosomal pellet obtained from HEK293T cells treated with control sgRNA (sgCtrl) or SERBP1 sgRNA (sgSERBP1) along with INK128 treatment for 48 hours. **K**. In vitro kinase assay using recombinant mTORC1 and purified SERBP1 as a substrate in the presence of ^32^P-γ-ATP. Asterisk indicates the auto-phosphorylation of a mTORC1 component. INK128 is used to inhibit mTOR. **L**. Model showing the TORC1-mediated regulation of dormant 80S ribosome formation and translation recovery upon refeeding via Stm1/SERBP1.

Knockdown of SERBP1 in HEK293T cells using siRNAs showed no significant change in polysome profile under normal conditions in which mTORC1 is active (Figures 6D, 6E, and S6C). However, SERBP1 depletion decreased 80S ribosomes and increased 40S and 60S subunits compared to control cells, upon mTOR inhibition by INK128 (Figures 6D, 6E, and S6C). Thus, similar to Stm1, SERBP1 forms dormant 80S ribosomes upon mTOR inhibition. To investigate the effect of SERBP1 on ribosome preservation, we deleted SERBP1 using CRISPR sgRNAs and analyzed ribosome levels. In proliferating cells, SERBP1 deletion had no effect on ribosome content compared to control cells (Figures 6F and S6D). Upon long-term inhibition (48 hours) of mTOR by INK128 treatment, SERBP1 deleted cells showed a significant reduction in ribosome content (Figures 6G and 6H) and a significant increase in ribosome poly-ubiquitylation (Figures 6I and 6J), compared to the control cells. Thus, SERBP1-mediated formation of dormant 80S ribosomes is important for the protection of ribosomes in human cells.

We examined whether SERBP1, like Stm1, is regulated by mTORC1. To investigate whether SERBP1 is directly phosphorylated by mTORC1, we performed an in vitro kinase assay with recombinant mTORC1 and SERBP1. mTORC1 phosphorylated SERBP1 in an INK128-sensitive manner, suggesting that SERBP1 is a mTORC1 substrate (Figure 6K). Furthermore, to identify mTOR-dependent phosphorylation sites on SERBP1, we performed a phosphoproteomic analysis of INK128-treated and -untreated HEK293 cells. Inhibition of mTOR caused a significant reduction in the phosphorylation of SERBP1 at serine 199 and threonine 226 (Figure S6E). We also observed a trend for reduced phosphorylation at serine 25, 197, and 234 residues (Figure S6E). The above results suggest that SERBP1 is phosphorylated by mTORC1. Thus, the regulation of ribosome dormancy via mTORC1-mediated phosphorylation of a ribosome preservation factor is evolutionarily conserved.

## Discussion

Our findings demonstrate that eukaryotic cells utilize Stm1/SERBP1 to inactivate and preserve ribosomes in response to TORC1 inhibition or nutrient deprivation. We further show that the nutrient sensor TORC1 controls Stm1/SERBP1 in an evolutionarily conserved manner. Under nutrient-sufficient conditions, TORC1 phosphorylates Stm1/SERBP1 to prevent it from forming a complex with the 80S ribosome. Upon TORC1 inhibition, dephosphorylated Stm1/SERBP1 forms non-translating, dormant 80S ribosomes that are protected from proteasome-mediated degradation (Figure 6L).

The accumulation of non-translating 80S ribosomes upon TORC1 inhibition or nutrient deprivation was previously reported but not well understood (Barbet *et al*., 1996; Gandin *et al*, 2014; Larsson *et al*., 2012; Muller *et al*, 2019). Based on conventional wisdom, one would expect an accumulation of free 40S and 60S subunits due to inhibition of translation initiation. Our study suggests that the accumulation of 80S ribosomes upon TORC1 inhibition is due to the formation of dormant 80S ribosomes by Stm1/SERBP1. In dormant 80S ribosomes, Stm1/SERBP1 occupies the mRNA tunnel, thereby preventing translation and ‘gluing’ the two subunits together (Anger *et al*., 2013; Jenner *et al*, 2012). In vitro translation experiments showed that Stm1 represses translation via its N-terminal region (Balagopal & Parker, 2011; Hayashi *et al*, 2018). We show that the N-terminal 120 residues of Stm1 are important for the formation of dormant 80S ribosomes upon TORC1 inhibition. We note that the recombinant Stm1 protein used in previous in vitro experiments was purified from *E. coli* and thus not in the inhibited, phosphorylated state. Thus, Stm1/SERBP1 prevents inappropriate translation initiation under TORC1-inhibited conditions, in addition to preserving non-translating ribosomes from proteasomal degradation (see below).

mTORC1 promotes translation at many levels. It stimulates ribosome biogenesis and tRNA synthesis (Mayer & Grummt, 2006). It promotes translation initiation by phosphorylating and inhibiting eIF4E-binding proteins (4EBPs), thereby allowing assembly of the eIF4F complex (Gingras *et al*, 2004; Proud, 2019). TORC1 also promotes translation initiation by phosphorylating initiation factor eIF4B (Ma & Blenis, 2009). Our data suggest that TORC1 also activates translation via phosphorylation and inhibition of Stm1. Thus, activation of dormant ribosomes via phosphorylation of Stm1 is yet another mechanism by which TORC1 promotes translation, underscoring the importance of translation as a readout of TORC1 signaling in the control of cell growth.

Our proposal that Stm1 forms dormant 80S ribosomes invokes the necessity of an additional mechanism to recycle 40S and 60S subunits during the resumption of translation. A previous study showed that glucose starvation, a condition where TORC1 is known to be inhibited, causes an accumulation of 80S ribosomes. During recovery from glucose starvation, Dom34 (duplication of multi-locus region 34) and Hbs1 (Hsp70 subfamily B suppressor 1) split vacant 80S ribosomes to recycle the two subunits (Pisareva *et al*, 2011; van den Elzen *et al*, 2014). Agreeing with our observation that Stm1 is required for the formation of dormant 80S ribosomes, in the absence of Stm1, Dom34 and Hbs1 were dispensable for resumption of translation (van den Elzen *et al*., 2014). Another study demonstrated that overexpression of Stm1 was toxic in strains lacking Dom34 (Balagopal & Parker, 2011), suggesting that Stm1 and Dom34 have opposing functions. However, a recent in vitro study concluded that Stm1-bound 80S ribosomes are resistant to splitting by Dom34-Hbs1 (Wells *et al*., 2020). Since this recent in vitro study lacked TORC1-mediated phosphorylation, we propose that TORC1-dependent Stm1 phosphorylation might be assisting in the recycling of dormant 80S ribosomes by Dom34 and Hbs1.

In yeast, upon TORC1 inhibition, ribosomes are considered to be degraded mainly via ribophagy (Kraft *et al*., 2008). However, we observed enhanced proteasome-mediated degradation of ribosomes in the absence of Stm1/SERBP1, with no effect on ribophagy. Furthermore, inhibition of proteasomal degradation upon rapamycin treatment partly prevented ribosome turnover, in both wild-type and *stm1*Δ cells. This suggests that under TORC1 inhibition, ribosomes are degraded by both proteasome and ribophagy (Kraft *et al*., 2008). A recent study in mammalian cells showed that proteasomal degradation, rather than ribophagy, is the major mechanism of ribosome degradation upon mTOR inhibition (An *et al*, 2020). How does Stm1/SERBP1 protect dormant 80S ribosomes from proteasomal degradation? One possible explanation is that the interface of the two ribosomal subunits, which is masked in the 80S ribosome, is recognized by ubiquitin ligases. It will be of interest to identify the mechanisms and the ubiquitin ligases that are involved in ubiquitylation and degradation of ribosomal subunits upon nutrient starvation.

TORC1 in yeast phosphorylates Stm1 to stimulate protein synthesis in response to nutrients, in vegetatively growing cells or germinating spores. mTORC1 similarly phosphorylates the Stm1 ortholog SERBP1 in mammalian cells. Although we did not investigate the physiological context in which mTORC1 regulates SERBP1, we speculate that this regulation could be important in the survival and activation of quiescent cells such as stem cells, immune cells, and gametes.

## Acknowledgements

We thank Prof. Michael W. Van Dyke (Kennesaw State University), Prof. Jonathan R. Warner (Albert Einstein Institute, NY) and Prof. Maurice S. Swanson (University of Florida) for antibodies against Stm1, Rpl3 and Pab1, respectively. We thank Dr. Nikolaus Dietz and Prof. Timm Maier for recombinant mTORC1. S.S. was supported by EMBO LTF 2016. M.H. acknowledges the Swiss National Science Foundation (grant number 179569 and NCCR RNA and Disease), the Louis Jeantet Foundation, and the Canton of Basel.

## Author Contribution

S.S., J.H., S.B., and D.R., conducted the experiments, S.S. and M.H. designed the experiments, analyzed the data, and wrote the manuscript.

## Method details

### Bacterial cells

DH5α *E. coli* cells were used to generate all the plasmid constructs used in the study. DH5α was grown in LB medium at 37°C.

### Yeast cells

Yeast strains used in this study are mainly generated using Tb50a background (Beck & Hall, 1999). Yeast cells are usually grown in YPD media at 30°C. For inhibition of TORC1, liquid cultures are treated with 200 nM rapamycin (prepared in 90 % ethanol 10 % Tween 20) or YPD agar plates containing 4 to 6 ng/mL rapamycin. For nitrogen starvation experiment, cells are grown in YPD to 0.6-0.8 OD_600_ and then washed once with sterile water and resuspended in synthetic media (containing yeast nitrogen base without ammonia and glucose) for the indicated time points. For restimulation of starved cells, 10X amino acid mix is used for the mentioned time points. Strains with deletion and chromosomal tags were generated in the background of TB50a (*MATa leu2-3,112 ura3-52 trp1 his3 rme1*) using homologous recombination. Oligonucleotides used in this study are listed in table S2.

### Human cell lines

HEK293T and HEK293 cells were obtained from ATCC. Cells were cultured in DMEM with high glucose supplemented with 10% Fetal bovine serum (FBS), glutamine, and Pen/Strep and incubated at 37°C, and 5% CO_2_. siRNA-mediated knock-down and sgRNA-mediated knock-outs were performed using JetPrime transfection reagent.

### Polysome profile analysis

The yeast cells are grown to 0.6 to 0.8 OD_600_ and treated with 100 μg/ml CHX for 2 min and harvested by centrifugation 2000g for 2 min at 4°C. The cell pellets are washed once with PBS containing CHX and once with the lysis buffer (20 mM Tris-HCl pH7.4, 50 mM KCl, 5 mM MgCl_2_, 1 % Triton X100, 1 mM DTT, 100 μg/mL CHX, 0.1 mM PMSF, 40 U/ mL RNasein plus) and resuspended in lysis buffer. Cells were lysed by bead beating method using vibrax for 15 min and the lysates were harvested. 100 μg of total RNA was loaded onto TH641 tubes containing 5-50% sucrose density gradient (in 20 mM Tris-HCl pH7.4, 150 mM KCl, 5 mM MgCl_2_, 1 mM DTT, 100 μg/mL CHX) and spun for 2 hours at 36000 rpm. For the low salt conditions, 50 mM KCl is used, while for high salt, 300 mM KCl is used. For analyses of ribosome content, 100 μg of cell extract was treated with 50 mM EDTA and then loaded onto 5-25% sucrose density gradient containing 50 mM KCl and spun for 4 hours at 36000 rpm using TH641 rotor (Sorvall). The cell lines were lysed by hypotonic lysis buffer (10 mM Tris-HCl pH7.4, 1.5 mM KCl, 5 mM MgCl_2_, 0.5 % Triton X100, 0.1 % Sodium deoxycholate, 1 mM DTT, 100 μg/mL CHX, EDTA free Protease Inhibitor, 40 U/ mL RNasein plus), incubated at 4°C for 15 min and the KCl concentration is adjusted to 100 mM prior to the centrifugation to obtain cell extract. The gradient conditions used were similar to the yeast lysates. After centrifugation, the gradients were analyzed at absorbance 260 nm using BIOCOMP fraction analyzer and 9 or 14 equal volume fractions were collected. For the immunoblot of the fractions, individual fractions were precipitated by 10% TCA, acetone washed twice and resuspended in 2X Laemmli buffer (36 % Glycerin, 15% 2-mercaptoethanol, 3.6% SDS, 90mM Tris pH 6.8, 0.07% Bromophenol Blue). For the mass spectrometric analysis, 10% TCA precipitated fractions were washed with acetone twice and resuspended in urea buffer compatible with mass spectrometry (see mass spec protocol). For the ribosome isolation using sucrose cushion, cell lysates were layered over 0.6 ml of 1M sucrose solution in TLA120.1 tube and centrifuged at 75000 rpm for 2 hours. The supernatant and the pellets were analyzed by immunoblot.

### Translation efficiency assay

For puromycin incorporation assay, yeast cells from specified conditions were treated with puromycin (100 μg/mL) for 15 min at 30°C. The reaction was stopped by 10% TCA (Trichloroacetic acid). The cells were washed with 95% acetone and resuspended in 0.2 M NaOH (with 0.1 mM PMSF) for 10 min at room temperature and lysed with 2x Laemmli buffer at 95°C for 15 min. The lysates were loaded onto 4-20% PAGE gels and immunoblotted for puromycin.

### Spore germination analysis

The diploid yeast cells were sporulated using sporulation media. The tetrads were dissected on YPD agar using a micromanipulator. The germination event was monitored microscopically after dissection and the number of spores achieving the 2-cell stage within 6 hours of dissection was counted. For each biological replicate, the germination of 14 spores was monitored. The plates were further incubated at 30°C to monitor colony formation. For the analyses of ribosome content in spores, RPL24 was GFP tagged on both chromosomes and the resultant cells were sporulated and monitored for the expression of GFP in tetrads by fluorescent microscopy.

### Immunoprecipitation

Yeast cells were resuspended in 50 mM Tris-HCl (pH 8.0), 100 mM KCl, 5 mM MgCl_2_, 1 mM PMSF, 1% Triton X100, 1x complete mini protease inhibitor, 1x PhosSTOP (hereafter referred to as IP buffer) and lysed using bead-beating. Lysates are harvested by centrifugation at 13000g for 10 min at 4°C to remove cell debris. Protein concentration in the supernatant was determined using BCA assay. For immunoprecipitation of GFP, GFP-Trap (ChromoTek) was used. 50 μl of magnetic beads was incubated with 2 mg of lysate for 3 hours at 4°C with gentle rotation. After 4 times washing with IP buffer without the detergent, the bound GFP tagged proteins were eluted with 2x Laemmli buffer and were resolved by SDS-PAGE for subsequent immunoblot analysis.

For purification of recombinant Stm1 and Stm1^AAA^, 0.6 OD_600_ cultures of BL21 strains containing either pGEX6P-GST-Stm1 or pGEX6P-GST-Stm1AAA were induced with 100 mM IPTG for 3 hours and lysed by sonication in lysis buffer containing PBS, 1% TritonX100 and 1 mM PMSF. Cell lysates were incubated with GST beads for 3 hours and washed 5 times with PBS. The beads were incubated with 1x Kinase assay buffer containing HRV 3C protease. The cleaved Stm1 was collected from the supernatant.

### In vitro Kinase assay

Recombinant SERBP1 (origene) or Stm1 or Stm1^AAA^ were incubated with 100 nM of purified mTORC1 (Timm Maier laboratory, Biozentrum) in a kinase assay buffer (50 mM HEPES pH 7.4, 10 mM MnCl_2_, 1 mM EGTA, 0.0025% Tween 20, 2.5 mM DTT, 10 μM ATP, 4 μCi ^32^P−γ-ATP) with DMSO or INK128 200 nM. The reaction is incubated at 30°C for 30 min. The reaction is stopped using 1x Laemmli buffer and boiled at 95°C for 10 min. The proteins were resolved using SDS-PAGE and either directly exposed to radiographic films or transferred to a membrane and then exposed to radiographic films.

### Analysis of ribosome composition

Yeast cells were grown to 0.5 OD_600_ and treated with 200 nM rapamycin for 1 hour. The preparation of cell extracts and polysome profile were performed as mentioned above. Each polysome profile was collected into 14 equal fractions and precipitated using TCA. The precipitates were acetone washed and resuspended in a resuspension buffer containing 8M Urea, 0.1M ammonium bicarbonate, and 5 mM TCEP. Samples were incubated at 37°C for 1h and treated with chloroacetamide at a final concentration of 15 mM. After an incubation of 30 min at 37°C, samples were diluted 8 times with 0.1M ammonium bicarbonate and sequencing-grade modified trypsin (1/50, w/w; Promega, Madison, Wisconsin) was added, and proteins were digested for 12 h at 37°C shaking at 300 rpm. Digests were acidified (pH<3) using TFA and desalted using iST cartridges (PreOmics, Martinsried, Germany) according to the manufacturer’s instructions. Peptides were dried under vacuum and stored at −20°C. Dried peptides were resuspended in 0.1% aqueous formic acid and subjected to LC–MS/MS analysis using a Orbitrap Fusion Lumos Mass Spectrometer. The acquired raw-files were imported into the Progenesis QI software. The generated mgf-file was searched using MASCOT against a Saccharomyces Cerevisiae database. The database search results were filtered using the ion score to set the false discovery rate (FDR) to 1% on the peptide and protein level, respectively, based on the number of reverse protein sequence hits in the dataset. Quantitative analysis results from label-free quantification were processed using the SafeQuant R package v.2.3.2. to obtain peptide relative abundances.

### Phosphoproteomic analysis

For yeast phosphoproteomics, cells were grown in YPD to 0.4 OD_600_ and treated with 200 nM rapamycin for 2 hours. For the phosphoproteomics of HEK293, 90% confluent cells were treated with DMSO or INK128 for 2 hours. The cells were lysed in 2M Guanidine HCl, 0.1M ammonium bicarbonate, 5 mM TCEP, and phosphatase inhibitors by bioruptor. Lysates were incubated for 10 min at 95°C and let cool down to room temperature followed by the addition of chloroacetamide at a final concentration of 15 mM. After an incubation of 30 min at 37°C, samples were diluted to a final Guanidine-HCl concentration of 0.4M using 0.1M ammonium bicarbonate buffer. Proteins were digested by incubation with trypsin (1/100, w/w) for 12 h at 37°C. After acidification using 5% TFA, peptides were desalted using C18 reverse-phase spin columns according to the manufacturer’s instructions, dried under vacuum, and stored at −20°C until further use. Peptide samples were enriched for phosphorylated peptides using Fe(III)-IMAC cartridges on an AssayMAP Bravo platform as recently described (Post *et al*, 2017). Phospho-enriched peptides were resuspended in 0.1% aqueous formic acid and subjected to LC–MS/MS analysis using an Orbitrap Fusion Lumos Mass Spectrometer. The acquired raw-files were imported into the Progenesis QI software. The generated mgf-file was searched using MASCOT against a human database or a Saccharomyces cerevisiae database. The database search results were filtered using the ion score to set the false discovery rate (FDR) to 1% on the peptide and protein level, respectively, based on the number of reverse protein sequence hits in the datasets. Quantitative analysis results from label-free quantification were processed using the SafeQuant R package v.2.3.2. (PMID:27345528, https://github.com/eahrne/SafeQuant/) to obtain peptide relative abundances. This analysis included global data normalization by equalizing the total peak/reporter areas across all LC-MS runs, data imputation using the knn algorithm, summation of peak areas per protein and LC-MS/MS run, followed by calculation of peptide abundance ratios.

For the analysis of phosphorylation on immune-precipitated Stm1 by mass-spectrometry, GFP-tagged Stm1 were pulled down from yeast cells with or without rapamycin treatment using GFP-Trap and subjected to on-bead digestion with Trypsin (Soulard *et al*, 2010). The peptides were purified using C18 columns and analyzed by mass spectrometry.

---

**Figure S1.**
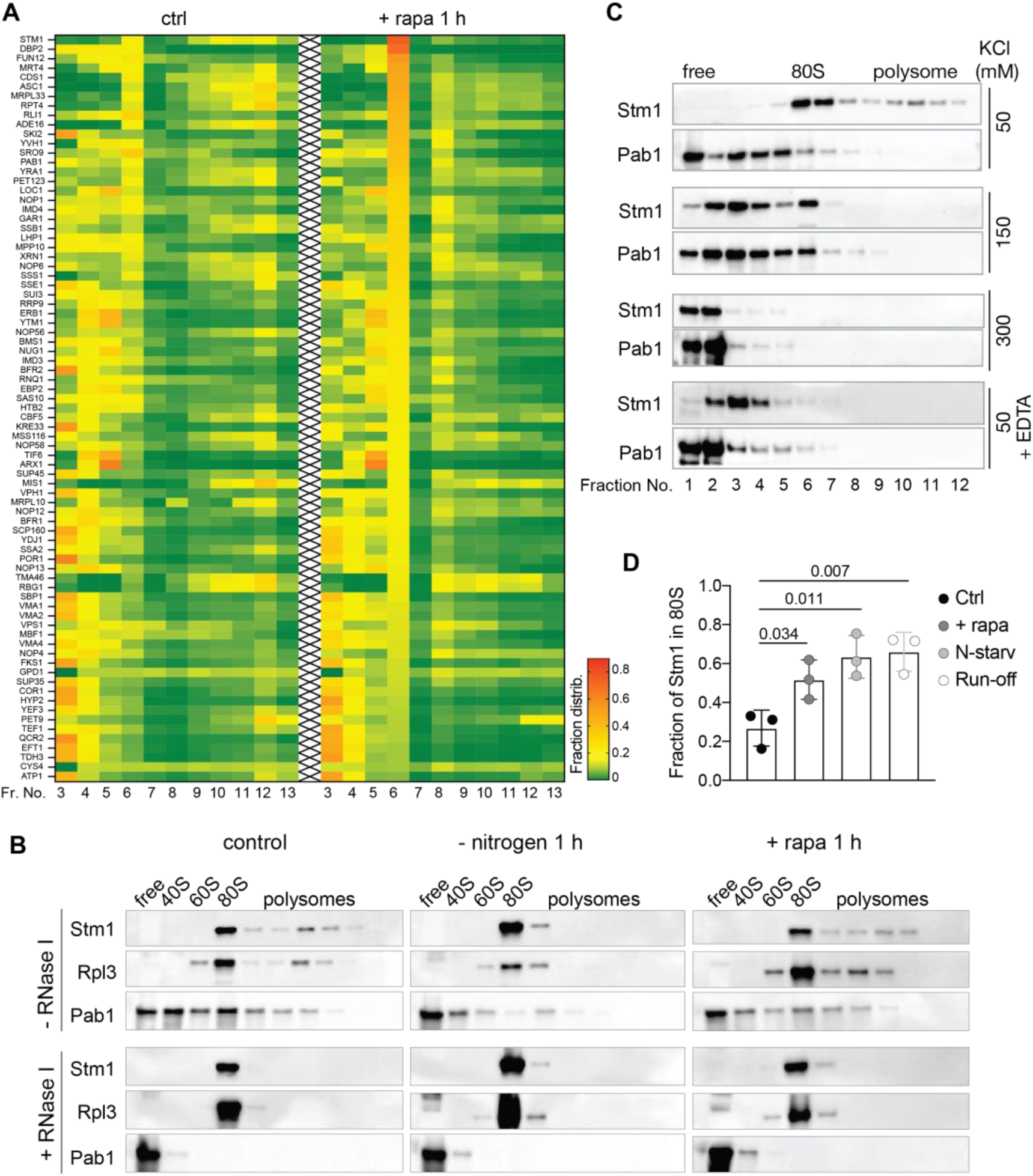
Characterization of the interaction of Stm1 with ribosomes. **A.** Heat map showing the fraction distribution of most abundant (top 80) ribosome-associated proteins and their response to rapamycin treatment. **B.** Immunoblots of Stm1, Rpl3, and Pab1 in fractions obtained from polysome profiles of yeast cells under control condition, nitrogen starvation for 1 hour, or rapamycin treatment for 1 hour. The polysome profiles were analyzed with or without RNase I treatment for 10 min using 50 mM KCl containing 5-50% sucrose density gradients. Representative blots from a minimum of three independent experiments are shown. **C.** Immunoblot of Stm1 and Pab1 in the polysome profile fractions obtained from sucrose density gradients containing 50, 150, or 300 mM KCl, or from cell extracts treated with 50 mM EDTA (analyzed in 50 mM KCl sucrose density gradients). **D.** Relative fraction distribution of Stm1 in 80S fraction. Minimum of three replicates are analyzed by multiple t-tests.

**Figure S2.**
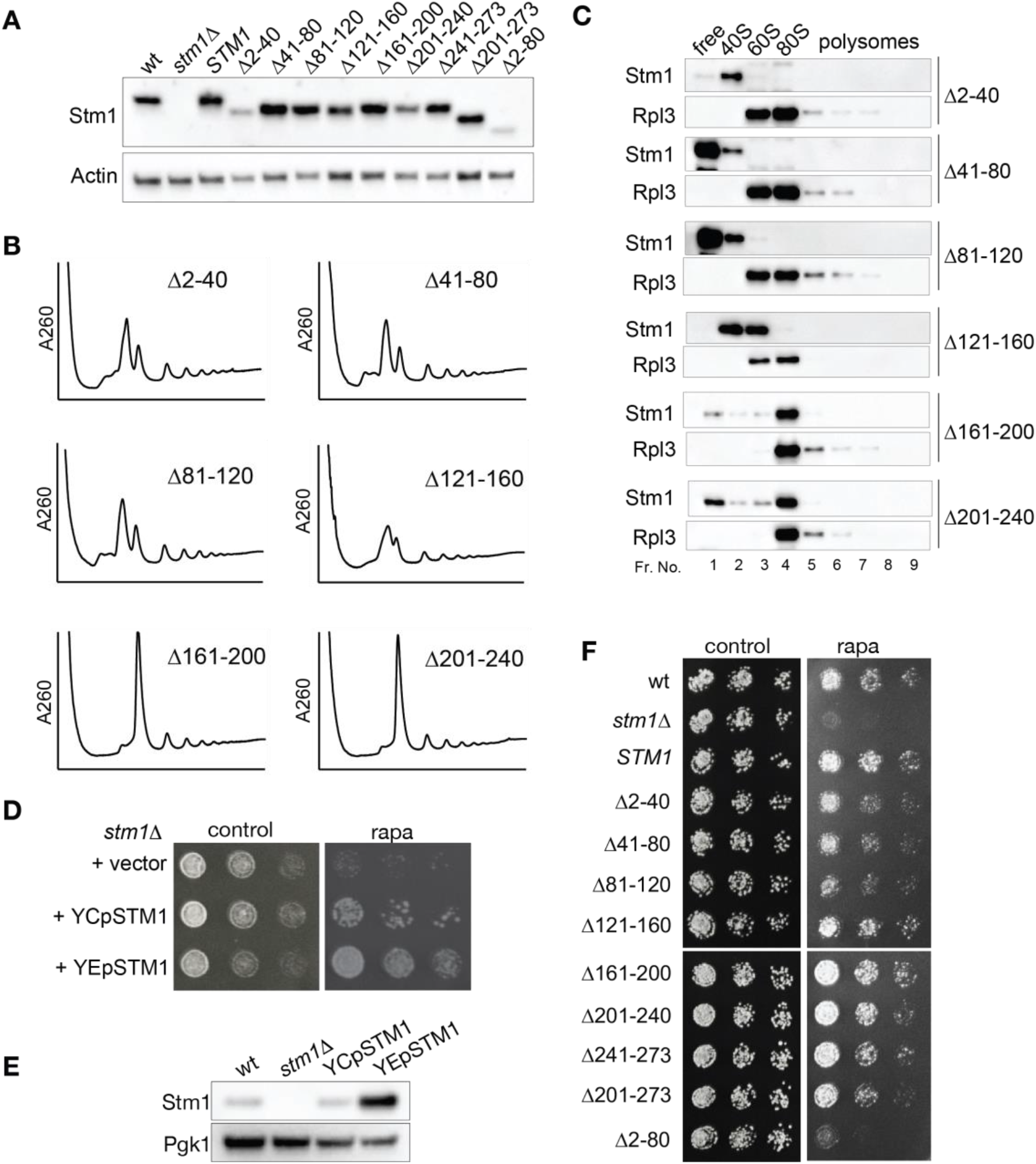
Stm1 forms vacant 80S ribosomes upon TORC1 inhibition. **A.** Immunoblot of Stm1 and Actin in *stm1*Δ cells harboring either vector alone, wild-type, or various deletion mutants of Stm1 in nutrient-sufficient conditions. **B.** Polysome profiles of *stm1*Δ cells containing various deletion mutants of Stm1 upon nitrogen starvation for 1 hour. **C.** Immunoblot of Stm1 across the polysome profile fractions shown in Figure S2B. 9 equal volume fractions of each polysome profile were precipitated and analyzed by immunoblot for Stm1 and Rpl3. At least two biological replicates for each mutant were analyzed. **D.** Growth of *stm1*Δ cells expressing either vector alone, Stm1 to the wild-type level (YCp*STM1,* single-copy plasmid) or overexpression of Stm1 (YEp*STM1,* multi-copy plasmid) on YPD agar plates with or without rapamycin. **E.** Immunoblot of Stm1 in *stm1*Δ, wild-type and Stm1-overexpressed cells. **F.** Growth of *stm1*Δ cells containing various deletion mutants of Stm1 on YPD agar plates with or without rapamycin.

**Figure S3.**
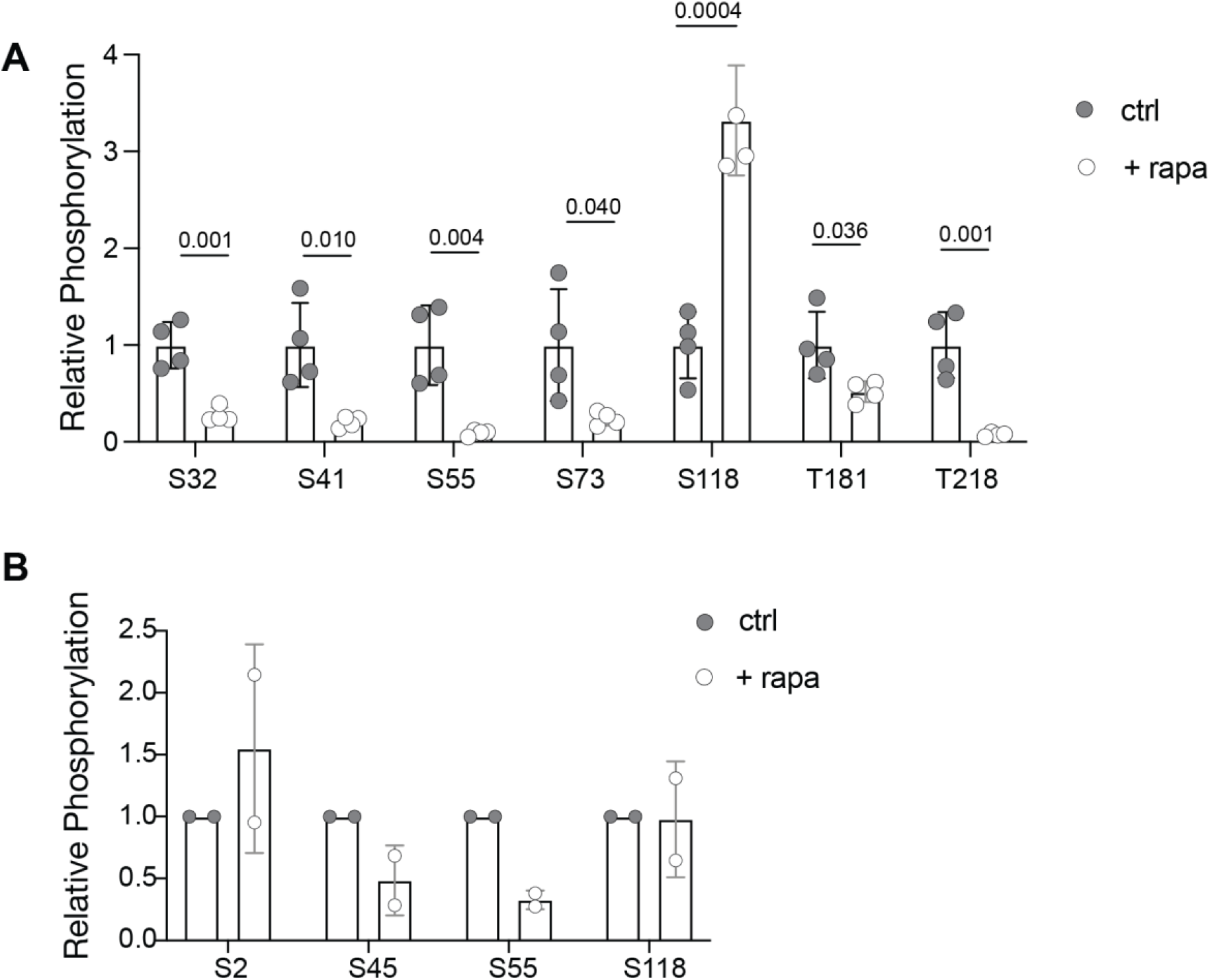
TORC1 phosphorylates Stm1. **A.** Relative phosphorylation of Stm1 identified from phosphoproteomic analysis of yeast cells treated with or without rapamycin for 2 hours. Four biological replicates are used and analyzed by multiple t-tests. **B.** Relative phosphorylation of Stm1 identified by mass spectrometry. Stm1-GFP was pulled down from yeast cells treated with or without rapamycin for 1 hour and subjected to mass spectrometric analysis. Two biological replicates were used.

**Figure S4.**
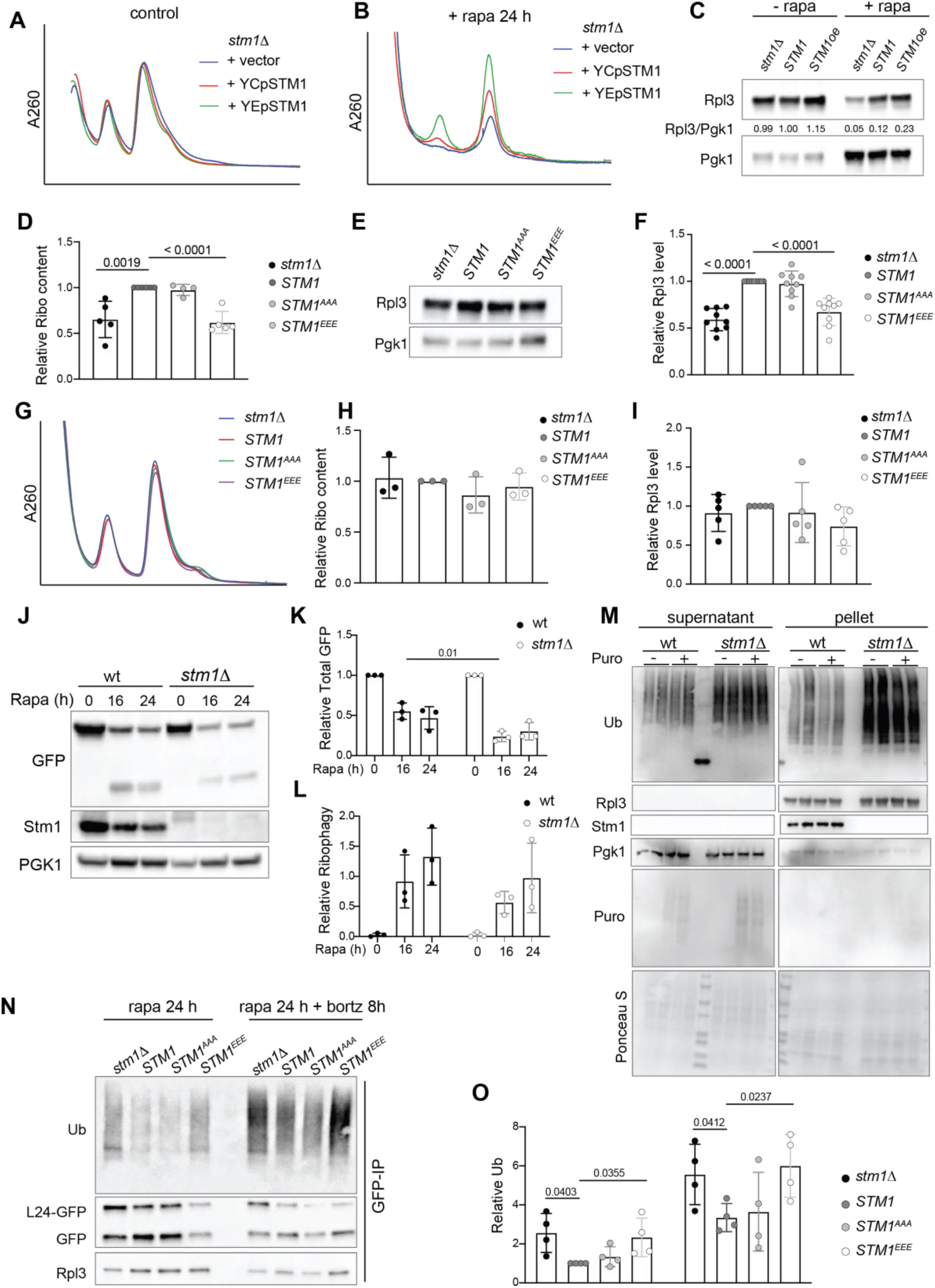
Stm1 preserves ribosomes under TORC1 inhibited conditions. **A-B**. Ribosomal subunit profiles of *stm1*Δ cells containing no (+ vector), wild-type (+ YCp*STM1*), or overexpressed (+ YEp*STM1*) levels of Stm1 without (**A**) or with (**B**) rapamycin treatment for 24 hours. For the ribosomal subunit profile analyses, the cell extracts were treated with 50 mM EDTA and separated on 5-25% sucrose density gradients. **C.** Immunoblot of Rpl3 in *stm1*Δ cells containing no (+ vector), wild-type (+ YCp*STM1*), or overexpressed (+ YEp*STM1*) levels of Stm1 along with or without rapamycin treatment for 24 hours. **D.** Ribosome content in *stm1*Δ, wild-type, *STM1^AAA^*, and *STM1^EEE^*cells starved for nitrogen for 24 hours, measured from ribosomal subunit profiles. Minimum of five replicates are used for P-value calculation. **E-F**. Immunoblot (**E**) and the quantification (**F**) of Rpl3 in *stm1*Δ, wild-type, *STM1^AAA^*, and *STM1^EEE^* cells under nitrogen starvation for 24 hours. At least six biological replicates are subjected to statistical analysis using multiple t-tests. **G-H.** Ribosomal subunit profiles (**G**) and quantification of relative ribosome content (**H**) of *stm1*Δ, wild-type, *STM1^AAA^*, and *STM1^EEE^* cells under nutrient-sufficient conditions. **I**. Quantification of Rpl3 in *stm1*Δ, wild-type, *STM1^AAA^*, and *STM1^EEE^* cells under nutrient-sufficient conditions. **J-L**. Immunoblot of Rpl24-GFP and free GFP (**J**), quantification of total GFP relative to Pgk1 (**K**), and free GFP relative to Rpl24-GFP (**L**) in wild-type and *stm1*Δ cells treated with rapamycin for 16 and 24 hours. At least 3 replicates are used for multiple t-test analysis. **M.** Immunoblots for ubiquitylation in supernatant and ribosomal pellet obtained from wild-type and *stm1*Δ cells starved for nitrogen (24 hours) along with or without puromycin treatment. Immunoblots for Stm1, Rpl3, Pgk1, and puromycin are also shown. **N-O**. Immunoblot for ubiquitin (**N**) and its quantification (**O**) after pull-down of Rpl24-GFP in *stm1*Δ, wild-type, *STM1^AAA^*, and *STM1^EEE^* cells treated with rapamycin for 24 hours along with or without bortezomib for 8 hours (in *pdr5*Δ background). Minimum of four replicates are quantified and analyzed using multiple t-tests.

**Figure S5.**
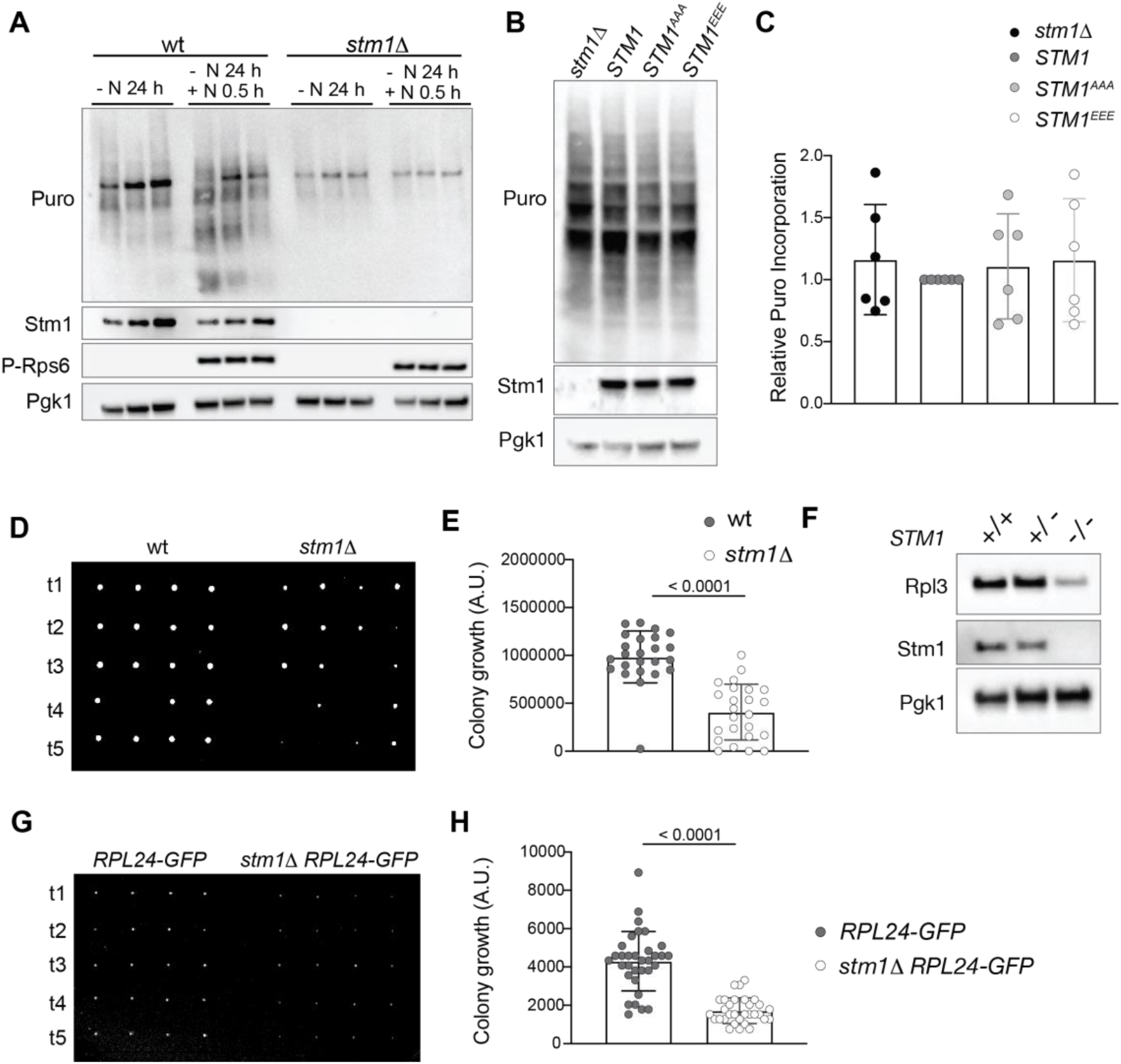
Stm1 is required for translation recovery. **A**. Puromycin incorporation in wild-type and *stm1*Δ cells subjected to nitrogen starvation for 24 hours and restimulated with amino acids for 30 minutes. Immunoblots for phospho-Rps6, Stm1, and Pgk1 are shown. Three biological replicates are shown for each condition. **B-C**. Puromycin incorporation (**B**) and its quantification (**C**) in *stm1*Δ, wild-type, *STM1^AAA^*, and *STM1^EEE^* cells under nutrient-sufficient conditions. Immunoblots for puromycin, Stm1 and Pgk1 are shown. Six biological replicates are analyzed by multiple t-tests. **D-E**. Growth of germinated spores (**D**) and its quantification (**E**) upon dissection of tetrads obtained from diploid wild-type and *stm1*Δ/*stm1*Δ strains on YPD agar media after 48 hours of incubation. (Each row shows 4 spores from individual tetrads of wild-type and *stm1*Δ which are labeled as t1, t2, etc.) **F**. Immunoblots for Rpl3, Stm1, and Pgk1 in diploid wild-type (TB50a), heterozygous *stm1*Δ, and homozygous *stm1*Δ strains grown in sporulation media. **G-H**. Growth of germinated spores (**G**) and its quantification (**H**) upon dissection of tetrads obtained from diploid control (*RPL24-GFP*) and *stm1*Δ *RPL24-GFP* cells on YPD agar media after 30 hours of incubation.

**Figure S6.**
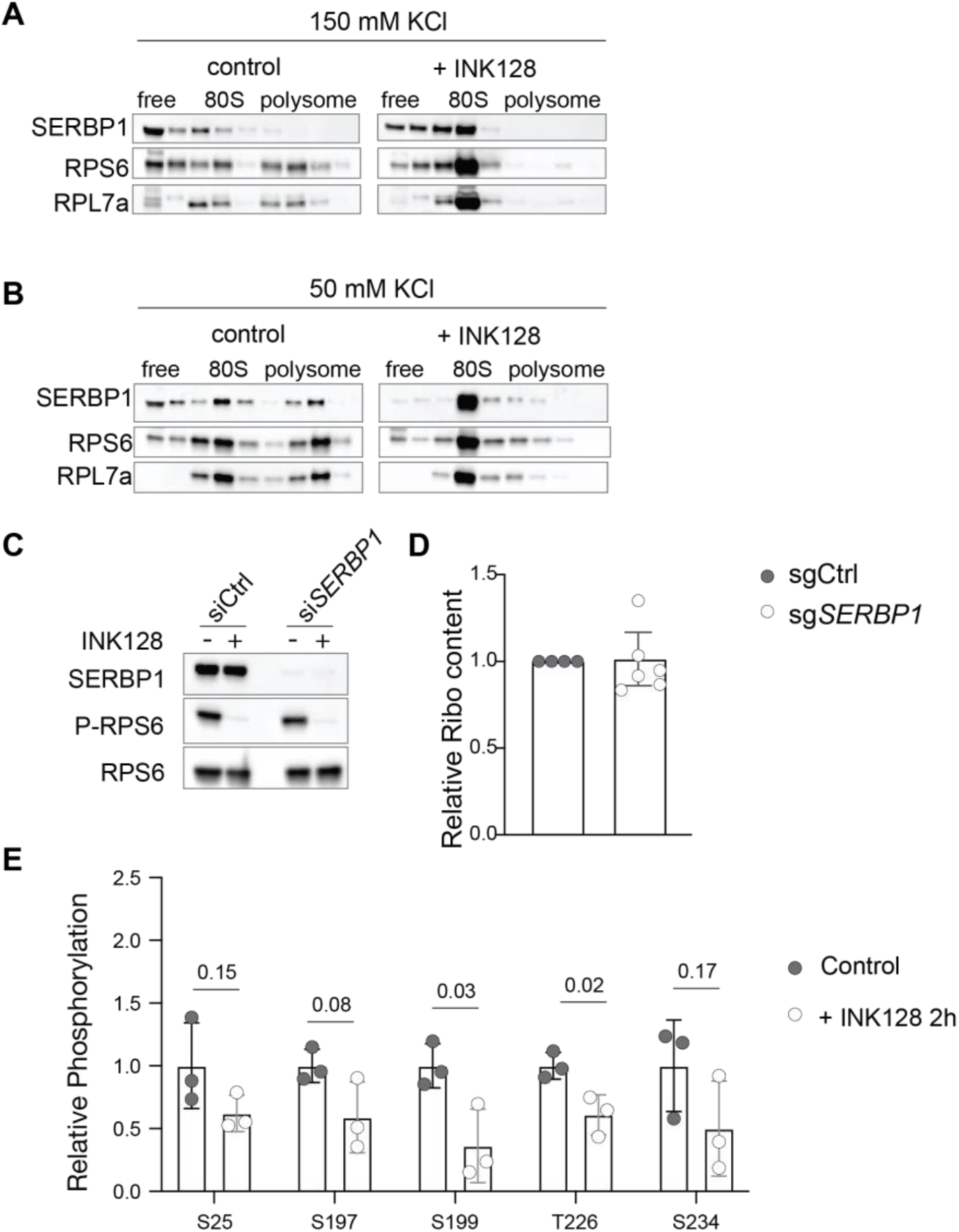
SERBP1 preserves ribosomes in human cells. **A-B**. Immunoblots of SERBP1, RPS6, and RPL7a across the polysome fractions of HEK293T cells treated with DMSO or INK128 for 1 hour. Polysome profiles were analyzed on 150 mM KCl (**A**) or 50 mM KCl (**B**) containing sucrose density gradients. **C**. Immunoblots of SERBP1, phospho-RPS6 (S240/244), and RPS6 in HEK293T cells treated with control-or SERBP1-siRNA, with or without INK128 for 1 hour. **D**. Quantification of ribosome subunit analysis of HEK293T cells treated with control sgRNA (sgCtrl) or SERBP1 sgRNA (sgSERBP1) under normal conditions (Related to Figure 6F). Minimum of four replicates are analyzed for each condition. **E**. Relative phosphorylation of SERBP1 identified from phosphoproteomic analysis of HEK293 cells treated with or without INK128 for 2 hours.

